# Revealing a Correlation between Structure and *in vitro* Activity of mRNA Lipid Nanoparticles

**DOI:** 10.1101/2024.09.01.610730

**Authors:** Xiaoxia Chen, Jialin Fang, Xue Ge, Mengrong Li, Fan Jiang, Liang Hong, Zhuo Liu

## Abstract

mRNA-loaded lipid nanoparticles (mRNA-LNPs) represent a promising platform for disease prevention, cancer immunotherapy, and gene editing, etc. Despite the success of mRNA-LNPs based vaccines, the relationship between their structure and efficacy remains elusive. Here, we generated a series of mRNA-LNPs with varied structural properties and *in vitro* cellular activities by altering their processing and storage conditions to investigate the structure-activity relationship (SAR). Our findings revealed a moderate anticorrelation between particle size distribution and *in vitro* activity. Importantly, the intensity of a characteristic peak, as detected by small angle X-ray scattering (SAXS), demonstrated a strong correlation with *in vitro* activity while the encapsulation efficiency is high. Additionally, the peak width and area were moderately correlated and anticorrelated with activity, respectively. These observations suggest that a more ordered internal structure is likely associated with enhanced *in vitro* activity of mRNA-LNPs. Further analysis using ^31^P nuclear magnetic resonance indicated that lyophilization might induce phase separation of mRNA and lipids within the LNPs, leading to a diminished SAXS peak and reduced *in vitro* activity. Overall, our study establishes an SAR for mRNA-LNPs, highlighting that a more ordered internal structure correlates with higher efficacy, which could be instrumental in high-throughput screening of LNP libraries.

## Introduction

The emergency use authorizations of two mRNA-based COVID-19 vaccines in 2020 garnered significant attention from both the academic and industrial community.^1, 2^ These vaccines employ mRNA-containing lipid nanoparticles (mRNA-LNPs), where the mRNA encodes the viral spike protein, prompting an immune response.^3^ This vaccination strategy builds on the success of LNPs in gene therapy and lipophilic drug delivery, which utilize ionizable lipids, helper phospholipids, cholesterol, and polyethylene glycol-lipids (PEG-lipids).^4^ Beyond infectious diseases, mRNA-LNPs hold promise for cancer immunotherapy, protein replacement therapy, and *in vivo* gene editing.^5, 6^ In June 2024, the FDA approved Moderna’s mRNA-based respiratory syncytial virus vaccine, mRNA-1345, marking the third mRNA-LNP vaccine to reach the market, further validating the potential of mRNA-LNPs in immunotherapy.^7^

Despite these exciting successes in application, the structure-activity relationship (SAR) of mRNA-LNPs remains incompletely understood, hindering their rational design and further optimization. This gap highlights the need for an efficient approach to high-throughput screening of mRNA-LNP formulations based on nanoparticle structure. Researchers have attempted to bridge the gap between nanoparticle structure and transfection potency using advanced techniques such as cryogenic electron microscopy (cryo-EM),^8, 9^ small angle neutron scattering (SANS),^10, 11^ and small angle X-ray scattering (SAXS).^12–15^ Although cryo-EM and SANS offer high-resolution insights into morphology, lipid distribution, and internal structure, their complexity and time-consumption of sample preparation prevent them from being applied in high-throughput experiments.^9, 10, 16–20^ In contrast, synchrotron SAXS allows for structural analysis with simple sample preparation and rapid data collection, making it more practical for SAR investigations.^12–15^ Owing to only one or two peaks at ∼0.1 Å^-1^ can be distinguished while probing the mRNA-LNPs by SAXS,^10, 16, 18^ previous studies have primarily focused on core structures^12, 13, 15^ or introduced structure-forming helper lipids^12–14^ to enrich structural information. However, changes in the shell structure have also been reported to influence mRNA-LNP efficacy.^9–11, 16^ Additionally, the use of structure-forming helper lipids may obscure a direct understanding of the SAR. Therefore, there is a pressing need for *in situ* investigations of mRNA-LNP SAR without external interference to achieve a deeper understanding.

In this work, we prepared 58 mRNA-LNP samples with varying *in vitro* cellular activities by manipulating processing and storage conditions. To explore the relationship between structure and activity, we assessed key structural parameters, including encapsulation efficiency (EE%), hydrodynamic radius (R_h_), radius of gyration (R_g_), and the degree of internal structural order. The *in vitro* activity of each mRNA-LNP sample was quantified by using the luminescence intensity within the identical amount of mRNA during transfection. We observed a strong correlation between the intensity of a SAXS peak at ∼0.13 Å^-1^ and in *vitro* activity while encapsulation efficiency is above 0.88, with the peak width and area showing moderate positive and negative correlations, respectively. These findings suggest that an ordered internal structure, as indicated by SAXS, enhances mRNA-LNP efficacy. Further ^31^P Nuclear Magnetic Resonance (NMR) analysis confirmed that lyophilization might induce phase separation in the core of mRNA-LNPs, leading to a weakened SAXS peak and reduced activity. Additionally, particle size distribution exhibited a moderate anticorrelation with activity across all the 58 samples. Our results establish a structure-activity relationship (SAR) for mRNA-LNPs, indicating that a more ordered internal structure may contribute to higher efficacy. This insight could be valuable for high-throughput screening of LNP libraries for mRNA loading and for monitoring efficacy of certain lipids formulation during processing and storage.

## Results

### Preparing the mRNA-LNP samples with varied *in vitro* activity

In this study, we firstly utilized two different chips, herringbone and Tesla mixer, to prepare the mRNA-LNP samples. The mRNA used in our work, encoding an enhanced green fluorescent protein (eGFP) linked to firefly luciferase (fLuc) with length of 2,856 nucleotides (nt). The *in vitro* transfection efficacy of the mRNA-LNP samples was assessed in four different cells, including human skeletal muscle cells (HskMC), Hepatoma G2 (HepG2) cells, Henrietta Lacks cells (HeLa), and human embryonic kidney 293 cells transformed with sheared simian virus 40 large T antigen (HEK293T). We quantified fLuc expression by measuring luminescence intensity within the identical amount of mRNA in each sample to evaluate transfection potency.

As illustrated in Figure 1a and the expanded cryo-EM images in Figure S1 of the Supporting Information (SI), the formulation prepared using the herringbone mixer yields predominantly spherical nanoparticles. The LNPs were further characterized by SAXS, with the distinct peak at ∼0.1 Å□¹ (Figure S2) providing definitive evidence for the encapsulation of mRNA within the lipid matrix, thereby confirming their identity as mRNA-LNPs. In contrast, the samples prepared using the Tesla mixer display bleb morphology (Figure 1b). Importantly, mRNA-LNPs with bleb morphology exhibited higher *in vitro* transfection efficacy in four different cell lines (Figure 1c), consistent with previous report.^8^ Given our goal of elucidating structure–activity relationships to inform development of more efficacious LNP drug products, we therefore focused on blebbed formulations for subsequent analysis.

**Figure 1.**
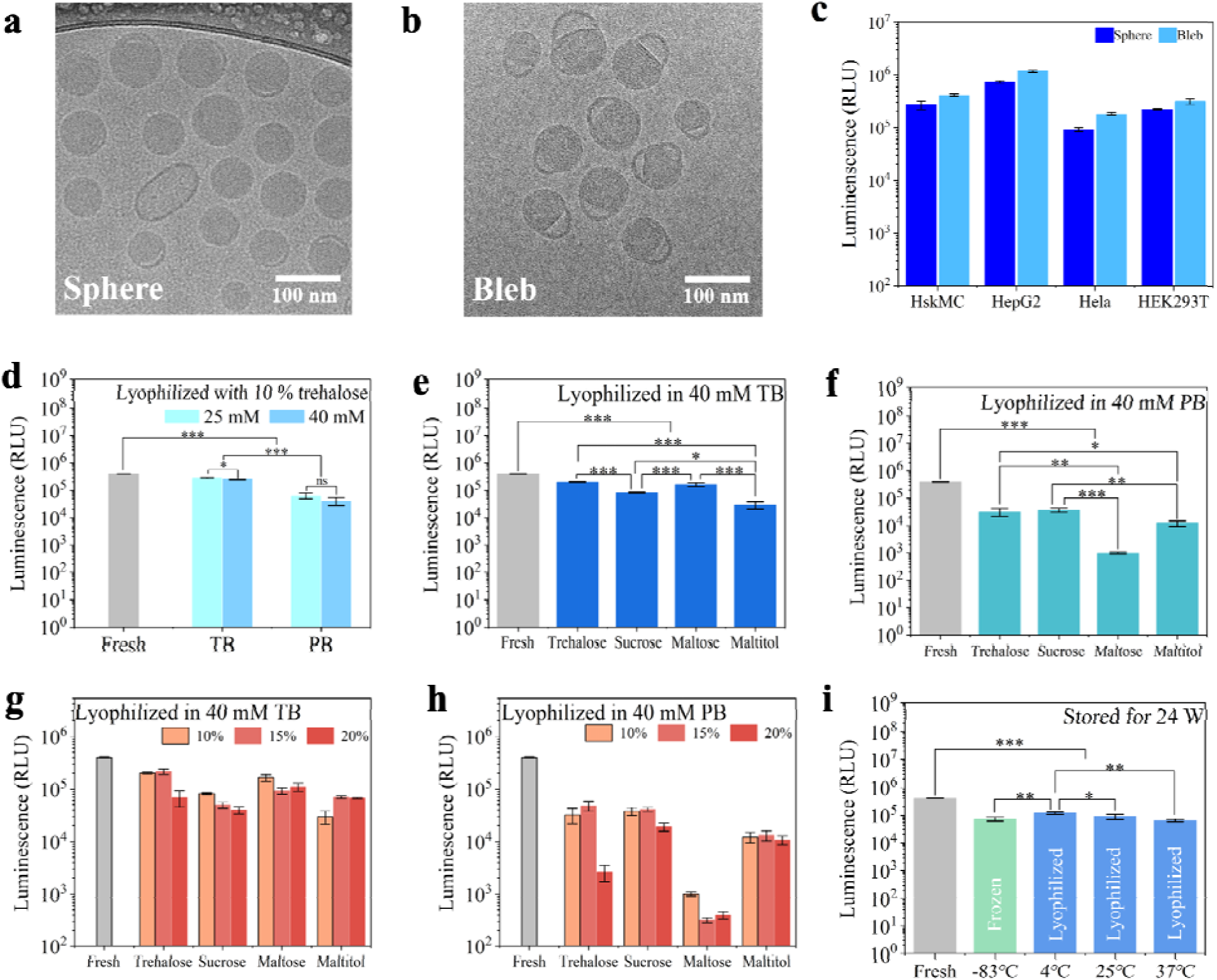
mRNA-LNP samples with varied *in vitro* activity. (a) Cryo-EM image of mRNA-LNPs predominantly exhibiting spherical structures. (b) Cryo-EM image of mRNA-LNPs primarily displaying bleb structures. (c) Luminescence of fresh spherical (dark blue) and blebbed (light blue) mRNA-LNPs in HskMC, HepG2, Hela, and HEK293T cells. (d) Luminescence of mRNA-LNPs lyophilized with 10% (w/v) trehalose in 25 mM Tris buffer (TB, cyan), 40 mM TB (blue), 25 mM phosphate buffer (PB, cyan), and 40 mM PB (blue) in HEK293T cells. (e) Luminescence of mRNA-LNPs lyophilized in 40 mM TB with 10% (w/v) lyoprotectants: trehalose, sucrose, maltose, or maltitol in HEK293T cells. (f) Luminescence of mRNA-LNPs lyophilized in 40 mM PB with 10% (w/v) lyoprotectants: trehalose, sucrose, maltose, or maltitol in HEK293T cells. (g, h) Luminescence of fresh mRNA-LNPs and mRNA-LNPs lyophilized with 10%, 15%, or 20% (w/v) lyoprotectants (trehalose, sucrose, maltose, or maltitol) in (g) 40 mM TB or (h) 40 mM PB in HEK293T cells. For 10% (w/v) lyoprotectants, the Lumi data are the same with those in panels e and f. (i) Luminescence of mRNA-LNPs frozen at –83 °C and stored for six months (6 M, green), compared with lyophilized mRNA-LNPs stored for 6 M at 4, 25, or 37 °C (blue). *, **, and *** indicate p ≤ 0.05, p ≤ 0.01, and p ≤ 0.001, respectively. ns in panel (d) denotes not significant. The fresh mRNA-LNP sample in panels d-i (gray) is the same as the blebbed fresh sample transfected in HEK293T cells in panel c.

To elucidate the SAR of mRNA-LNPs, it is essential to generate a series of mRNA-LNPs with diverse structural characteristics and *in vitro* cellular efficacy. Previous studies have demonstrated that processing and storage conditions significantly impact both the structure and efficacy of mRNA-LNPs.^17, 21, 22^ Accordingly, we prepared 58 distinct mRNA-LNP samples by varying their processing and storage conditions. Detailed methods for sample preparation are provided in the Materials and Methods, and a comprehensive list of sample information is available in Table S1. The samples were subjected to freezing or lyophilization under different buffer compositions, lyoprotectants, and storage conditions, while maintaining an identical chemical composition. Given that lyophilization can induce changes in structure and activity,^17, 21^ we anticipated sampling a broader structural spectrum. As depicted in Figure 1d-i, variations in sample preparation methods led to distinct luminescence (Lumi) in HEK293T cells. This wide distribution of Lumi values provided a robust dataset for examining the factors influencing the *in vitro* activity of mRNA-LNPs.

Figure 1d shows that mRNA-LNPs lyophilized with 10% (w/v) trehalose in Tris buffer exhibited higher *in vitro* activity than those prepared in sodium phosphate buffer, consistent with recent findings.^22^ Moreover, trehalose showed superior or similar protecting efficacy compared to other three lyoprotectants in both Tris and phosphate buffer (Figure 1e and 1f). Buffer concentration (25–40 mM) had little impact on activity in either Tris or phosphate systems (Figure 1d), whereas lyoprotectant concentration played a critical role (Figures 1g and 1h). Both 10% and 15% (w/v) trehalose in Tris buffer preserved mRNA activity after lyophilization, with 15% often yielding equal or slightly higher luminescence. Therefore, 15% trehalose in Tris buffer was selected for all subsequent lyophilization experiments to ensure maximal structural and functional preservation. Remarkably, this optimized formulation preserved higher mRNA-LNP activity at week 24 at 4 °C compared to frozen storage at –83 °C (Figure 1i). Among the tested conditions, 4 °C was identified as the most favorable storage temperature for lyophilized samples. Meanwhile, to generate a library covering a range of structural and functional states for SAR analysis, we then varied storage temperature and duration as suboptimal conditions (Figure S3).

The long-term stability of mRNA-LNPs in the optimal lyophilized condition have also been tested in HEK293T cells (see Figure S3). The size of lyophilized mRNA-LNPs stored at 4, 25, and 37 □ is ∼230 nm in 24 weeks, whereas the size of frozen mRNA-LNPs stored at -83 □ is ∼130 nm in the storage period. Nevertheless, both size distribution and encapsulation efficiency of lyophilized mRNA-LNPs stored at 4, 25, and 37 □ are comparable with those of frozen mRNA-LNPs stored at -83 □ in the storage period. More importantly, *in vitro* activity of lyophilized mRNA-LNPs stored at 4 □ can be comparable to that of frozen mRNA-LNPs stored -83 from 12 to 24 weeks, despite non-inferior activity of the frozen sample was observed from 0 to 12 weeks. Therefore, we have developed an optimal process of lyophilization for long-term storage of mRNA-LNPs, which can maintain comparable *in vitro* activity of mRNA-LNPs at 4 for 24 weeks with those stored at -83 □in the frozen state.

Collectively, these results underscore that maximizing the efficacy of lyophilized mRNA-LNPs requires careful selection of buffer species, lyoprotectant type and concentration, and storage conditions. It is important to note, however, that the lyophilization protocol established here is optimized for the specific lipid composition and mRNA length used in this study. For long-term storage of other mRNA-LNP formulations, re-optimization of buffer, lyoprotectant, and storage parameters will be necessary.^21–25^

To validate our results, we further examined the *in vitro* activity of mRNA-LNPs in three additional cell lines—HskMC, HepG2, and HeLa—alongside HEK293T cells. For each cell line, we tested eight representative mRNA-LNP formulations, including both frozen and lyophilized samples prepared with 15% (w/v) trehalose or 15% (w/v) sucrose in either Tris or phosphate buffer. As shown in Figure S4, across all four cell lines, mRNA-LNPs formulated in Tris buffer consistently demonstrated higher or comparable efficacy relative to those in phosphate buffer under the same storage conditions. Moreover, lyophilized samples exhibited lower luminescence intensity compared to their corresponding frozen counterparts in every cell line. The reproducibility of these trends across diverse cell types confirms the robustness of our results initially observed in HEK293T cells.

### Basic characterizations for the mRNA-LNP samples

To investigate the relationship between the structure of mRNA-LNPs and their *in vitro* activity, we first analyzed the particle size distribution. Using SAXS and dynamic light scattering (DLS), we observed that both the hydrodynamic radius (R_h_) and the radius of gyration (R_g_) of mRNA-LNPs exhibited minimal correlation with *in vitro* activity (Figure 2a and 2c). Also, the encapsulation efficiency of mRNA-LNPs is weakly correlated with *in vitro* activity (Figure 2d).

**Figure 2.**
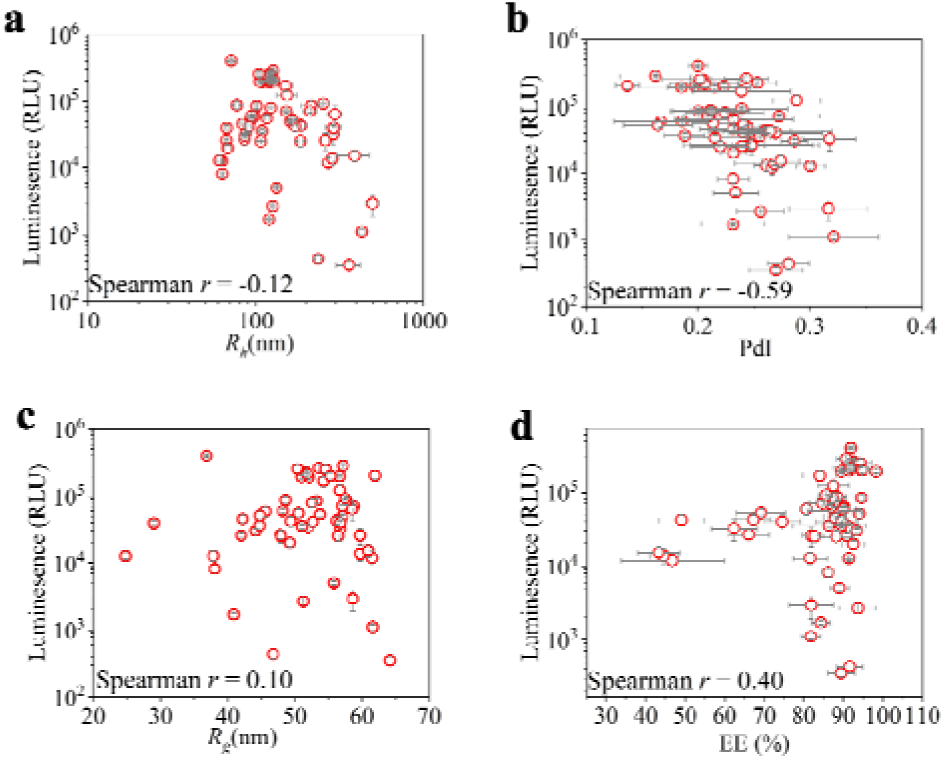
Basic characterizations for the mRNA-LNP samples. Luminescence of mRNA-LNPs plotted as a function of their (a) hydrodynamic radius (R_h_) and (b) polydispersity index (PdI), characterized using DLS. (c) Luminescence of mRNA-LNPs as a function of radius of gyration (Rg) characterized by SAXS. (d) Luminescence of mRNA-LNPs as a function of their encapsulation efficiency (EE%). All the samples were transfected in HEK293T cells.

However, the polydispersity index (PdI), reflecting particle size distribution, showed a moderate inverse correlation with protein expression in HEK293T cells (Figure 2b).

### Structural characterizations for the mRNA-LNP samples by SAXS

Given the lack of significant correlation between above physicochemical properties and *in vitro* activity, we extended our analysis to the internal structure of mRNA-LNPs, which has been reported to influence their delivery efficiency.^9, 11, 12, 14, 15, 26^ Notably, mRNA-LNPs with ordered internal structures, as revealed by SAXS, have been associated with enhanced intracellular mRNA delivery compared to those with disordered cores.^9, 12, 14, 26^ To examine these structural variations, we employed SAXS to analyze both mRNA-free LNPs and mRNA-LNPs.

Figure 3a shows a distinct peak at ∼0.13 Å^-1^ in the SAXS curve of mRNA-LNPs, indicating an ordered structure, whereas this peak is absent in mRNA-free LNPs. The SAXS data for the mRNA-free LNPs were fitted using a generic broad peak model constituted by a Porod curve and a Lorentz peak (Figure 3b, Table S2). For the mRNA-LNPs, the SAXS curve was fitted with a linear combination of this broad peak model, with the peak position fixed from the mRNA-free LNP fitting, and an additional Gaussian peak, allowing all parameters to vary (Figure 3c and 3d). Detailed methods and results for the data fitting are provided in Materials and Methods and the SI (Figure S5, Tables S2 and S3). This approach enabled us to resolve multiple structural features, including the intensity, width, and area of the Gaussian peak, the area of the Lorentz peak derived from the broad peak model for mRNA-free LNPs, and the ratio of Gaussian to Lorentz peak areas.

**Figure 3.**
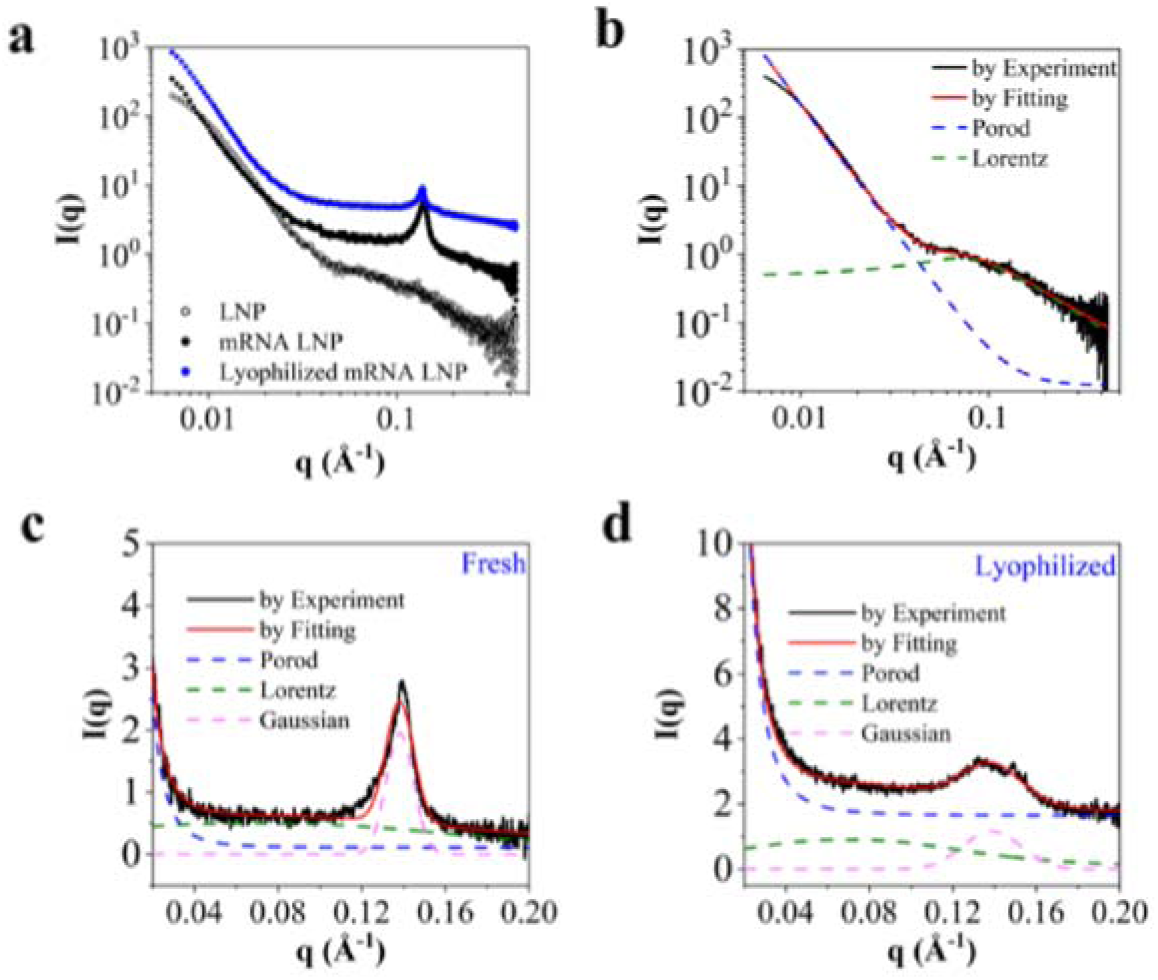
Structural characterizations for mRNA-free LNPs and mRNA-LNPs. (a) SAXS profiles of mRNA-free LNPs (black circles), fresh mRNA-LNPs (black dots), and lyophilized mRNA-LNPs (blue dots). (b) SAXS data for mRNA-free LNPs were fitted using a broad peak model. (c) SAXS data for fresh mRNA-LNPs were fitted with a broad peak model and a Gaussian peak. (d) SAXS data for lyophilized mRNA-LNPs with 15% (w/v) trehalose, stored at 25 °C for 12 weeks, were also fitted using the same model. In panels b-d, the blue, green, and pink dashed lines represent the Porod scaling law, Lorentz peak, and Gaussian peak, respectively, while the solid black and red lines correspond to experimental and fitting results.

### Correlation between structure and *in vitro* activity of mRNA-LNPs

The morphology of mRNA-LNPs was characterized by capturing metrics such as hydrodynamic radius (R_h_), polydispersity index (PdI), and cryo-EM images. SAXS further provided insights into structural features, including R_g_ and the subsequent fitting parameters of the peak at ∼0.13 Å^-1^. To identify key structural parameters influencing mRNA-LNP efficacy, we first performed principal component analysis (PCA). Dimension 1 of the PCA accounted for 35.9% of the total variance (Figure 4b) and was dominated by particle size and internal structural features, particularly the SAXS peak intensity at ∼0.13 Å□¹ (Figure 4a). More importantly, we observed that the SAXS peak intensity at ∼0.13 Å□¹ is close with luminescence at both direction and intensity in the biplot of PCA (Figure 4c). This analysis suggested that the ordered internal structure, as indicated by this peak, plays a central role in determining *in vitro* activity.

**Figure 4.**
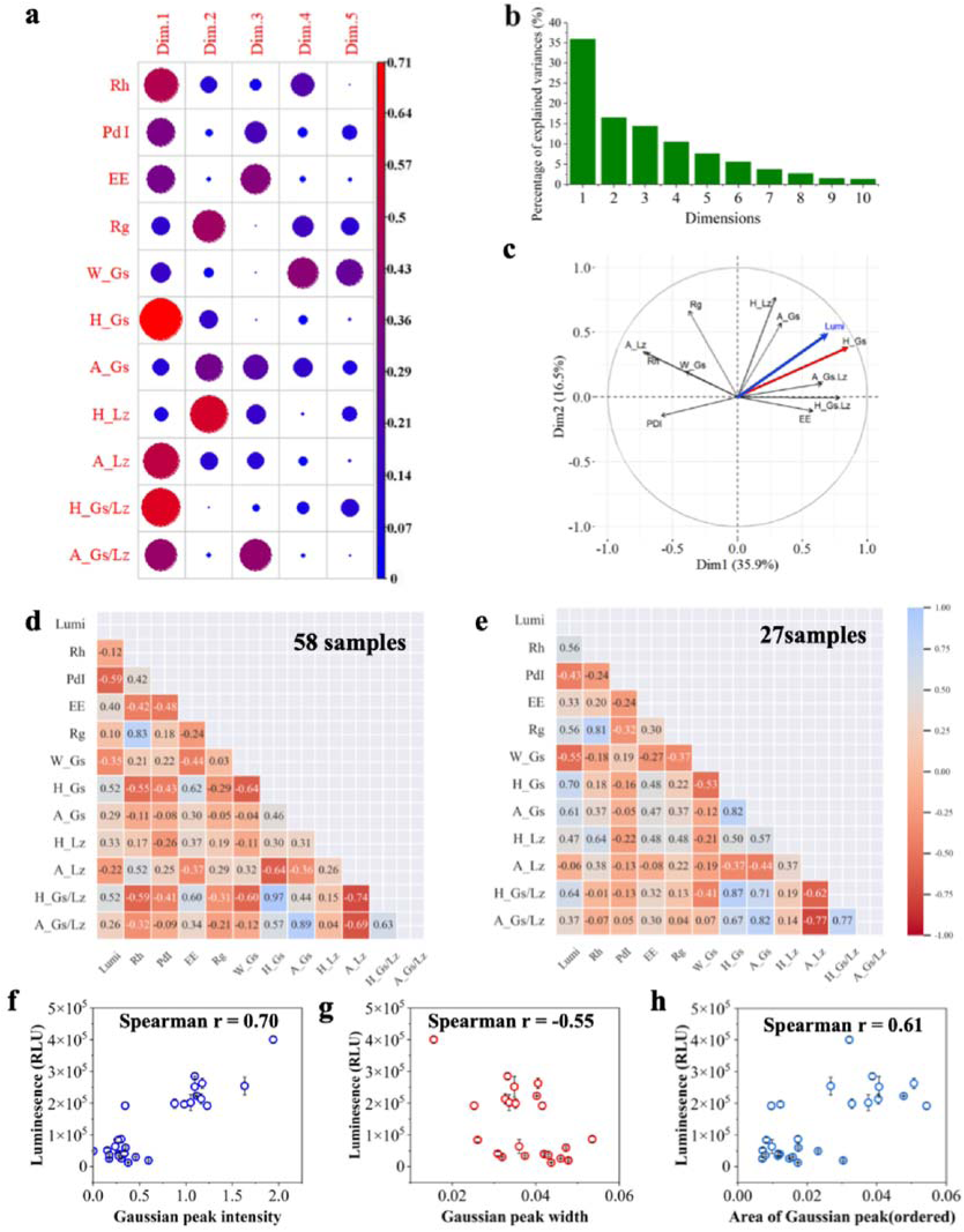
Correlation between structural parameters and *in vitro* activity of mRNA-LNPs. Principal component analysis (PCA) of structural parameters and in vitro efficacy of mRNA-LNPs. (a) Percentage of variance explained by the first nine principal components. (b) Contributions of structural indices to principal components 1–5. (c) Biplot of PCA in dimension 1 (Dim1) and dimension 2 (Dim2), where the blue and red arrows represent luminescence (Lumi) and the SAXS peak intensity at ∼0.13 Å□¹ (H_Gs), respectively. Color-coded heat maps depict the Spearman correlation coefficients between structural indices and in vitro activity for (d) 58 mRNA-LNP samples and (e) 27 mRNA-LNP samples with encapsulation efficiency greater than 0.88, with indices ranked according to their correlation strength. The parameters R_h_, PdI, EE, and R_g_ denote hydrodynamic radius, polydispersity index, encapsulation efficiency, and radius of gyration, respectively. W_Gs, H_Gs, and A_Gs correspond to the width, intensity, and area of the Gaussian peak derived from fitting the SAXS feature at ∼0.13 Å□¹, while A_Lz represents the area of the Lorentzian peak. The *in vitro* activity is expressed as Lumi, referring to the luminescence intensity of expressed firefly luciferase (fLuc). (f-h) Correlation of Lumi (for samples with encapsulation efficiency > 0.88) with (f) Gaussian peak intensity, (g) Gaussian peak width, and (h) Gaussian peak area derived from SAXS fitting.

To further assess structure–activity correlations, we employed Spearman’s rank correlation analysis across all 58 mRNA-LNP samples. As shown in Figure 4d, this analysis revealed a moderate negative correlation between polydispersity index (PdI) and luminescence, and a moderate positive correlation between the SAXS peak intensity at ∼0.13 Å□¹ and luminescence. Note that the SAXS peak at ∼0.13 Å^-1^ is much less pronounced if mRNA is separated with lipids (Figure 3a). To reveal the correlation between internal structure of mRNA-LNPs with their *in vitro* activity, it is also necessary to ensure most mRNA molecules are incorporated in the lipids. Thus, we refined our analysis by selecting a subset of 27 mRNA-LNP samples with encapsulation efficiencies above 0.88. Within this group, the Spearman correlation coefficient between SAXS peak intensity and luminescence reached 0.70, indicating a strong positive correlation (Figure 4e and 4f). Also, the peak height ratio of Gaussian peak and Lorentz peak have a Spearman correlation coefficient of 0.64 with the luminescence in the subgroup. In addition, the width and area of the Gaussian peak show moderate negative and positive correlations with activity, respectively (Figure 4g and 4h). These findings suggest that mRNA-LNPs with higher peak intensity, narrower width, and larger area—indicative of a more ordered internal structure—are more likely to exhibit enhanced *in vitro* efficacy.^18^ Conversely, the area of the Lorentz peak, associated with disordered structures, shows minimal correlation with activity (Figures 4). This implies that *in vitro* activity is largely independent of disordered structures within mRNA-LNPs. Besides, PdI exhibits a moderate inverse correlation with *in vitro* activity (Figure 4e), consistent with the trends observed across all 58 samples (Figure 4d).

To examine our results, we compared a freshly prepared mRNA-LNP sample with a representative lyophilized mRNA-LNP sample, focusing on encapsulation efficiency, structural characteristics, and *in vitro* activity. As illustrated in Figure 5a, both samples displayed comparable encapsulation efficiency, R_g_, and PdI, while the lyophilized sample exhibited a larger R_h_ compared to the fresh sample. Previous research has suggested that larger mRNA-LNPs can maintain or enhance transfection potency;^11, 27^ however, the lyophilized sample demonstrated reduced *in vitro* efficacy relative to the fresh one (Figure 5b). Both fresh and lyophilized mRNA-LNPs exhibited peaks at ∼0.13 Å^-1^ in SAXS characterization (Figure 5c). It is worth noting that the Gaussian peak derived from the SAXS curve of the fresh sample had greater intensity and area than that of the lyophilized sample, with similar peak widths between the two (Figure 5d). This finding suggests that the internal structure of lyophilized mRNA-LNPs is less ordered than that of the fresh sample.

**Figure 5.**
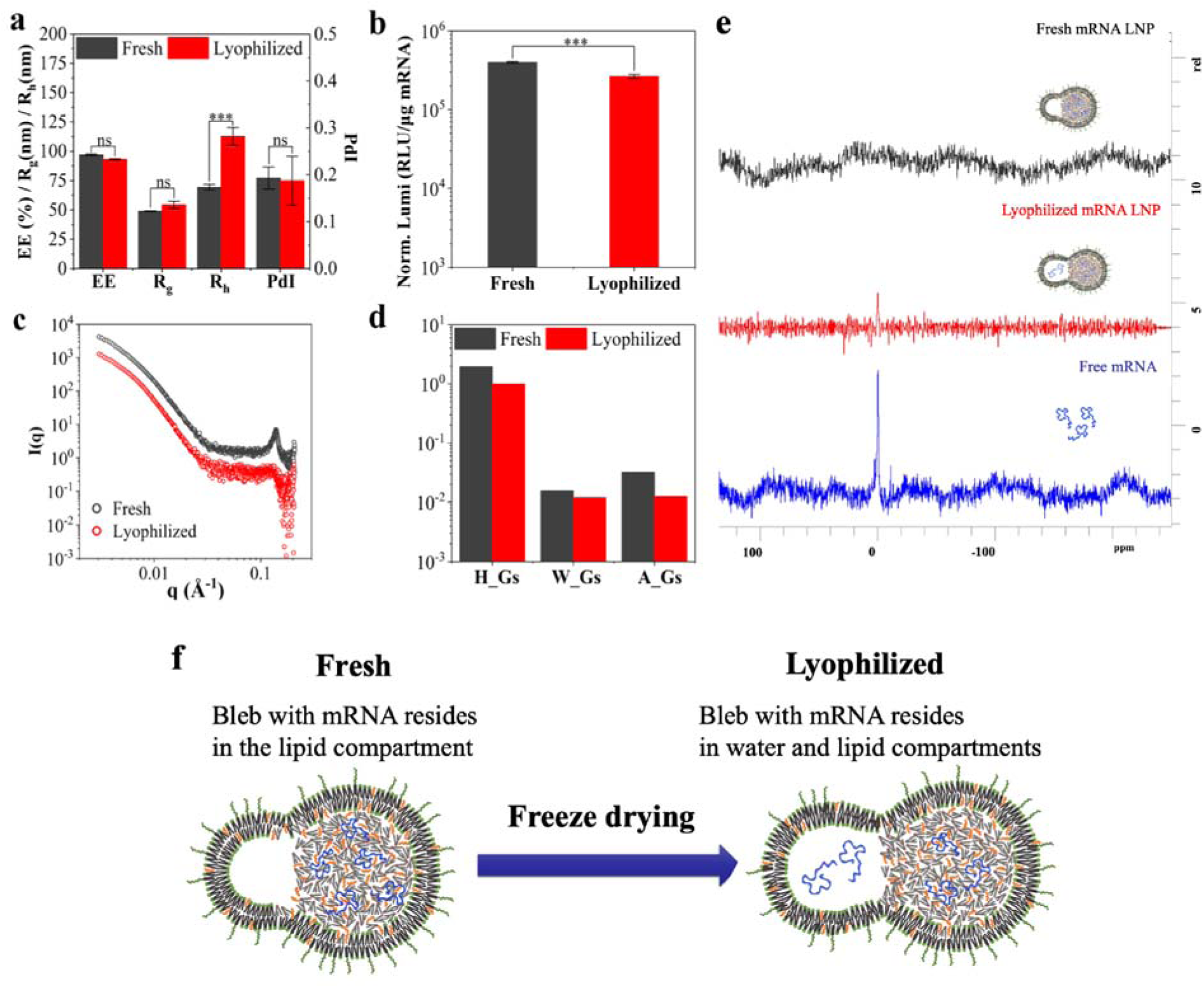
Comparative analysis of fresh and lyophilized mRNA-LNPs in the presence of 15% (w/v) trehalose in TB buffer. (a) A comparative evaluation of encapsulation efficiency (EE%), R_g_, R_h_, and PdI between fresh (black bars) and lyophilized (red bars) mRNA-LNPs. (b) A comparison of luminescence intensity (Luminescence) between fresh (black bar) and lyophilized (red bar) mRNA-LNPs. (c) SAXS curves for fresh (black dots) and lyophilized (red dots) mRNA-LNPs, highlighting structural differences. (d) A comparison of the intensities, widths, and areas of Gaussian peaks derived from fitting the SAXS data of fresh (black bars) and lyophilized (red bars) mRNA-LNPs. (e) ^31^P NMR spectra of free mRNA, fresh mRNA-LNPs, and lyophilized mRNA-LNPs, with upper, middle, and lower insets illustrating schematic representations of the structural organization of fresh mRNA-LNPs, lyophilized mRNA-LNPs, and free mRNA. (f) Schematic diagram for the probable impact of freeze drying on fresh mRNA-LNPs. In these schematics, blue lines represent mRNA molecules, cyan dots indicate water molecules, and other symbols denote lipid components.

To further explore mRNA distribution within the LNPs, we employed ^31^P nuclear magnetic resonance (NMR) spectroscopy. It is known that siRNA complexed with cationic lipids within LNPs displays broad “solid-state” ^31^P NMR resonances due to restricted mobility and large chemical shift anisotropy of the phosphate phosphorus.^28, 29^ In contrast, freely tumbling siRNA in an aqueous environment produces a narrow, easily detectable ^31^P NMR spectrum. We expected that fresh mRNA-LNPs would show no characteristic peak in the ^31^P NMR spectrum, while the lyophilized sample would resemble mRNA in an aqueous solution. Consistent with this expectation, Figure 5e shows a peak at approximately -0.6 ppm in the lyophilized mRNA-LNPs, similar to that of mRNA in solution, whereas the fresh mRNA-LNPs exhibited no peak. These observations suggest that lyophilization may cause partial segregation of mRNA molecules from lipids within the LNPs, leading to a bleb morphology with distinct water and lipid compartments, as depicted in Figure 5f. This phase separation likely results in a less ordered internal structure, as reflected by the reduced intensity of the SAXS peak at ∼0.13 Å^-1^ (Figure 5c). Given the strong correlation between the intensity of the SAXS peak at ∼0.13 Å^-1^ and the *in vitro* activity of mRNA-LNPs (Figure 4), it appears that a more ordered internal structure is associated with higher efficacy. This conclusion aligns with prior studies on mRNA-LNPs^12, 14, 15^ and is further supported by research on antisense oligonucleotide-loaded LNPs, which demonstrated that an ordered oligonucleotide-lipid phase in LNPs enhances endosomal escape and efficacy.^30^ Thus, the intensity of the characteristic SAXS peak attributed to the periodic structure of mRNA and lipids may serve as a reliable indicator for optimizing mRNA-LNP formulations.

## Discussion

Despite the recognized importance of the structure of mRNA-LNPs in determining bioactivity, the SAR remains poorly understood, largely due to limited resources and the complexity of structural characterization. For oligonucleotide-loaded lipid nanoparticles, Hammel et al. have identified the the disordered, H_II_, and Lα phases in the ASO-loaded LNPs by resolving the interlayer distance in cryo-EM and multiple peaks in SAXS to address the correlation between the internal structure and gene silencing activity.^30^ However, SAXS typically resolves only one or two peaks around ∼0.1 Å□¹ when probing mRNA-LNPs,^10, 16, 18^ and cryo-EM is unable to distinguish the internal structural characteristics for mRNA-LNPs. Hence, prior studies have primarily concentrated on core structures^12, 13, 15^ or relied on structure-forming helper lipids^12–14^ to enrich the available structural information of mRNA-LNPs. In parallel, several reports have also demonstrated that changes in the shell structure can modulate mRNA-LNP efficacy.^9–11, 16^ Moreover, the use of helper lipids may complicate the direct interpretation of structure–activity relationships. These obstacles hinder the accurate prediction of bioactivity of mRNA-LNPs.

To address this challenge, we prepared 58 mRNA-LNP samples with varying structures and *in vitro* activities through freezing and freeze-drying. Structural characterization was performed using SAXS, resolving a characteristic peak at ∼0.13 Å^-1^ in SAXS analysis. Furthermore, we used principal component analysis (PCA) to identify the key structural parameters influencing mRNA-LNP efficacy, and recognized that internal structural features, particularly the SAXS peak intensity at ∼0.13 Å□¹, plays a central role in determining *in vitro* activity. To further assess structure–activity correlations, we employed Spearman’s rank correlation analysis across all 58 mRNA-LNP samples. It is noteworthy that the intensity of the Gaussian peak derived from SAXS data showed a strong correlation with *in vitro* activity if the encapsulation efficiency is higher than 0.88, while the area and width of the Gaussian peak were moderately correlated and anticorrelated with activity, respectively (Figure 4). This finding offers a reliable index—the intensity of the characteristic Gaussian peak—for predicting mRNA-LNPs’ bioactivity and facilitates high-throughput screening of LNP formulations using SAXS.

It is noteworthy that this SAR was developed based on findings from a specific formulation composition, thereby we should be cautious when extrapolating this correlation to other LNP formulations. Interestingly, a recent study analyzed eight samples across four different LNP formulations and reported a strong positive correlation (Spearman r > 0.6) between the intensity of this very SAXS feature and *in vitro* activity in T cells.^31^ This independent confirmation in a multi-formulation context supports the robustness and general applicability of the structural descriptor we have identified. It reinforces that our SAR findings extend beyond the specific library space of our initial study and point toward a fundamental structural principle governing mRNA-LNP efficacy. Notably, this study only examined eight samples across four formulations, and concluded that SAXS peak intensity and encapsulation efficiency positively correlate with T-cell transfection efficiency. While this work further underscores the importance of nanoscale structure, the dataset is relatively limited and does not systematically probe a continuous spectrum of structural states. In contrast, our study investigates 58 mRNA-LNP samples with controlled variation in structure and performance, enabling a more robust quantitative assessment of structure–activity relationships. Using this expanded dataset, we find that encapsulation efficiency exhibits only a weak correlation with *in vitro* activity across the full sample set, and a strong correlation between SAXS peak intensity (∼0.13 Å□¹) and activity emerges only when encapsulation efficiency exceeds 0.88. Hence, our experimental results refine and contextualize the conclusions drawn from smaller-scale studies.

In addition to developing a methodology to correlate the SAR of mRNA-LNPs, we sought to understand the mechanisms underlying the loss of efficacy in mRNA-LNPs following lyophilization. Previous studies have indicated that the SAXS peak at ∼0.1 Å^-1^ is prominent only when mRNA is encapsulated within LNPs.^10, 19^ We propose that this SAXS peak originates from the periodic repeating distances between structural units formed by mRNA and adjacent ionizable lipids, consistent with prior SAXS^10^ and SANS^16^ findings. The high intensity, narrow width, and large area of the SAXS peak at ∼0.13 Å^-1^ suggest an ordered periodic distance of approximately 5 nm, indicating a more ordered internal structure, which is likely associated with higher *in vitro* activity (see Figure 4). After lyophilization and reconstitution, the mRNA-LNPs may adopt a bleb morphology characterized by water compartments containing mRNA molecules separate from lipid compartments, similar to the morphology observed in self-amplifying mRNA-LNPs.^19^ This separation likely results in a disordered internal structure, evidenced by a lower intensity Gaussian peak at ∼0.13 Å^-1^ in SAXS analysis, which correlates with diminished *in vitro* activity.

## Conclusion

In this study, we prepared 58 mRNA-LNPs with varied structures and *in vitro* activities by changing their storage conditions. Structural characteristics, including R_g_, R_h_, PdI, and the features of the SAXS peak at ∼0.13 Å^-1^, were determined using DLS and SAXS techniques. Concurrently, we measured the luminescence of expressed fLuc in HEK293T cells. Our findings indicate that the luminescence (Lumi) exhibits minimal correlation with encapsulation efficiency, suggesting that encapsulation efficiency is not directly associated with the *in vitro* activity of mRNA-LNPs. However, PdI, reflecting particle size distribution, shows a moderate negative correlation with *in vitro* activity, suggesting that a broad size distribution may reduce efficacy.

Additionally, the SAXS peak at ∼0.13 Å^-1^ was deconvoluted into two components: a Lorentz peak and a Gaussian peak, corresponding to contributions from disordered and ordered internal structures, respectively. We identified a strong correlation between the intensity of the Gaussian peak and the *in vitro* activity of mRNA-LNPs with encapsulation efficiency above 0.88, indicating that the intensity of the characteristic SAXS peak can serve as a predictive marker for *in vitro* activity under various conditions. Moreover, the area and width of the Gaussian peak displayed moderate positive and negative correlations with activity, respectively. Based on these observations, we propose that ordered internal structures formed by mRNA and adjacent ionizable lipids may enhance the *in vitro* activity of mRNA-LNPs. Further analysis using ^31^P NMR spectra revealed that lyophilization may cause separation of mRNA molecules from lipids within LNPs, leading to reduced Gaussian peak intensity and decreased efficacy. This finding underscores the impact of lyophilization on the stability and activity of mRNA-LNPs.

Overall, our study establishes a correlation between structure and *in vitro* activity of mRNA-LNPs using SAXS and introduces a methodology that can be applied to characterize LNPs throughout preparation, storage, and delivery stages. This workflow can be extended to identify critical formulation parameters within the vast formulation space, ultimately guiding the optimization of LNP performance.

## Materials and Methods

### Materials

The enhanced green fluorescent protein-linked firefly luciferase (eGFP-fLuc) encoding mRNA, with a length of 2,856 nucleotides, was procured from Yaohaibio Co., Ltd. (Jiangsu, China). This mRNA was synthesized via *in vitro* transcription, featuring a Cap1 structure at the 5’ end, pseudouridine substitution, and a 150-nucleotide poly(A) tail, with a purity exceeding 95%. The cationic ionizable lipid (CIL), heptadecan-9-yl 8-((2-hydroxyethyl)(6-oxo-6-(undecyloxy)hexyl)amino)octanoate) (SM-102, CAS: 2089251-47-6), was obtained from Sinopeg Biotechnology Co., Ltd. (Fujian, China). The 1,2-Distearoyl-sn-glycero-3-phosphocholine (DSPC, CAS: 816-94-4) was sourced from Merck (Darmstadt, Germany). Cholesterol (Chol, CAS: 57-88-5) and 1,2-Dilauryl-rac-glycero-3-methoxypolyethylene glycol 2000 (DMG-PEG2000, CAS: 160743-62-4) were supplied by Avanti Polar Lipids, Inc. (Alabaster, AL, USA). Lyoprotectants, including trehalose (Tre, CAS: 6138-23-4), sucrose (Suc, CAS: 57-50-1), maltose (Mal, CAS: 6363-53-7), and maltitol (Mai, CAS: 585-88-6), were purchased from Shaoxin Biotechnology Co., Ltd. (Shanghai, China).

### LNP formulation and preparation

The mRNA was dissolved in 100 mM citrate buffer (pH 4.0) at a concentration of 72 μg/mL, serving as the aqueous phase. Four lipids were dissolved in anhydrous ethanol at a molar ratio of CIL:Chol:DSPC:DMG-PEG2000 = 50:38.5:10:1.5. Both the aqueous and ethanol phases were equilibrated to room temperature prior to use. The mRNA-LNPs were prepared using a microfluidic device (NanoGenerator Flex, Precigenome LLC, CA, USA) at a total flow rate of 4 mL/min, with an aqueous-to-ethonal phase flow rate ratio of 3:1.

The mixture in the initial 5 seconds (∼0.25 mL) was discarded to eliminate the scrap attributed to non-equilibrium flow rates and residual buffer in the outlet channel. Most LNPs were formulated using the Tesla chip to generate blebbed nanoparticles, except for one mRNA-LNP sample with predominantly spherical morphology that was produced using the herringbone chip. This spherical formulation was served as a fresh spherical sample and was used for initial characterization. The primary mRNA-LNP suspension, containing 8 mM lipid and an N:P ratio of 6:1, was immediately diluted tenfold in Tris or phosphate buffer (pH 7.4) to reduce ethanol concentration and replace the buffer. Ultra-filtration at 3000 rpm using a 100 kDa MWCO Millipore filter was then employed to remove excess buffer. This buffer replacement was repeated to ensure that ethanol concentration in the final samples was reduced to less than 0.5%, while the mRNA-LNP was concentrated back to its initial volume prior to buffer replacement. For subsequent analyses, the mRNA-LNPs were further concentrated by ultra-filtration as needed. As a negative control, mRNA-free LNPs were prepared using 100 mM sodium citrate buffer (pH 4.0) as the aqueous phase, following the same method. A 30 kDa MWCO Millipore filter was used for buffer replacement and particle concentration of the mRNA-free LNPs.

### Lyophilization protocols

Concentrated mRNA-LNP suspension and 50 (w/v) % lyoprotectant in Tris- or Phosphate-buffer at pH 7.4 were mixed to yield mRNA-LNP samples with mRNA and lyoprotectant at final concentration of 54 μg/ml and 10∼20 (w/v) %, respectively. Glass vials (2 ml) were filled with 0.5 mL of the above mRNA-LNP formulation, and then frozen at -83 □ ahead of lyophilization by laboratory freeze-dryer (Alpha 2-4 LSCplus, Martin Christ Gefriertrocknungsanlagen, Germany) equipped with a LyoCube4-8 chamber. Lyophilization programs were set manually, with program parameters including running time and vacuum degree during the warm-up, major and secondary drying stages. Warm-up phase contains 2 stages, during which temperature of shelf was lower than -35 □ contributed to pre-freezing of shelf at -83 □. In the first warm-up stage, vacuum was created from 1 atm to 0.3 mbar in 40 min. Thereafter, 20 min evacuation kept going until vacuum reached 0.1 mbar. Primary drying was conducted at 0.1 mbar, with temperature increasing from -35 □ to 20 □ programmatically during following 39 hours. At the secondary drying stage, the vacuum was pumped to 0.01 mbar and kept for 9 hours, during which the temperature increased from 20 □ to 25 □. Once lyophilized, the vials were finally stoppered with rubber lids and sealed by a cap with sealing ring. Each vial was reconstituted with 500 µL of DEPC-treated water prior to evaluating the physicochemical properties and in vitro activity of the lyophilized mRNA-LNPs. Additionally, lyophilized samples were stored at 4°C, 25°C, and 37°C for varying durations to assess storage stability, with aqueous backup aliquots kept at -83°C in corresponding lyoprotectants.

### Dynamic light scattering (DLS)

The hydrodynamic radius (R_h_) and polydispersity index (PdI) of the mRNA-LNPs were determined using the NanoBrook Omni multi-angle particle size and zeta potential analyzer (Brookhaven Instruments, NH, USA). Prior to analysis, 25 µL of mRNA-LNPs was diluted with 500 µL of DEPC-treated water and transferred to a 50 µL disposable plastic cuvette (Brookhaven). Measurements were performed at a scattering angle of 90° using a 640 nm laser wavelength, with a dust filter engaged. The correlator was configured for nanoparticle analysis, and the solvent was set to water with default parameters. The refractive index of the particles was adjusted to 1.45, as ref. 10. Samples were equilibrated at 25°C for 120 seconds before measurement. The apparent hydrodynamic diameter was computed using the Einstein-Stokes equation, and the results are presented as the radius of an equivalent spherical particle.

### Cryo-Electron Microscopy (Cryo-EM)

The morphology of mRNA-LNPs was examined using cryogenic electron microscopy (cryo-EM). Sample preparation was carried out using a Leica EM GP2 automatic plunge freezer (Wetzlar, Germany). Briefly, 3 µL of LNP suspension, with a lipid concentration ranging from 5 to 10 mg/mL, was applied to a plasma-cleaned lacey copper grid coated with a continuous carbon film. Excess sample was removed by blotting, ensuring no damage to the carbon layer. The grids were then stored in liquid nitrogen until imaging. Cryo-EM imaging was performed on a Talos F200C G2 Microscope (Thermo Fisher Scientific, MA, USA) operated at 200 kV, and equipped with a Schottky thermal field emission electron gun. During the imaging process, the sample grids were maintained at temperatures below −170 °C. Images were captured using a FEI Ceta (4 k × 4 k) camera, and data acquisition was facilitated using the EPU and Serial EM software.

### Encapsulation efficiency

The encapsulation efficiency (EE%) of mRNA-LNPs was assessed using the Quant-iT RiboGreen RNA Quantification Kit, following the manufacturer’s instructions and conducted in a black 96-well plate. Briefly, 1 µL of mRNA-LNP suspension was diluted in 100 µL of Tris-EDTA (TE) buffer for measurement of free mRNA, or in TE buffer containing 2% Triton X-100 for measurement of total mRNA. Triton X-100 disrupts the LNPs, releasing the encapsulated mRNA, which allows for its detection. Subsequently, 100 µL of a RiboGreen solution, diluted 2000-fold for free mRNA and 200-fold for total mRNA, was added to the respective wells. RiboGreen, a fluorescent dye, exhibits enhanced fluorescence upon binding to nucleic acids. Fluorescence intensity was measured at 525 nm with an excitation wavelength of 425 nm using a Spark multimode microplate reader (Tecan, Männedorf, Switzerland). Calibration curves were generated for mRNA concentrations ranging from 0 to 100 ng/mL and 0 to 2000 ng/mL, prepared by sequential dilutions and mixed with the RiboGreen solutions in the same plate as the samples. The concentrations of free and total mRNA were determined from these calibration curves and used to calculate the encapsulation efficiency (EE%) using the formula [100 - *C*_free_ / *C*_total_ × 100] %, where *C*_total_ represented the total mRNA concentration in the presence of 2% Triton X-100, and *C*_free_ was the concentration of free or unencapsulated mRNA.

### *In vitro* cell transfection

Human Embryonic Kidney 293 cells stably expressing the sheared Simian Virus 40 Large T Antigen (HEK293T) (obtained from the Chinese Academy of Sciences Shanghai Institutes for Biological Sciences Cell Resource Center, Shanghai, China) were cultured according to established protocols. The cells were maintained in high-glucose Dulbecco’s Modified Eagle Medium (DMEM) (Thermo Fisher Scientific, CA, USA) supplemented with 10% fetal bovine serum (Gibco, Australia) and 1% penicillin/streptomycin. Cultures were incubated at 37 °C in a 5% CO_2_ atmosphere. For transfection, cells were seeded at a density of 3×10^5^ cells/mL in 24-well plates, with 500 μL per well, and allowed to adhere and proliferate for 24 hours. Transfection was carried out by adding 20 μL of mRNA-LNP suspension dropwise to the culture medium in triplicate wells, followed by a further 24-hour incubation period. The dosage of mRNA before lyophilization is ∼1 μg for each sample. mRNA expression in HEK293T cells was assessed by measuring fLuc activity using the Firefly Luciferase Reporter Gene Assay Kit (RG006, Beyotime Biotechnology, Shanghai, China), following the manufacturer’s instructions. After aspirating the culture medium, 100 μL of cell lysis buffer was added to each well. The cell lysate was centrifuged at 10,000×g for 5 minutes, and 40 μL of the supernatant was combined with 100 μL of Firefly Luciferase Assay Reagent. Luminescence was measured using a SpectraMax iD5 multi-mode microplate reader (Becton, Dickinson and Company, NJ, USA). The relative luminescence units (RLU) were reported as luminescence (Lumi). HepG2 and Hela cells were cultured and transfected by using the same protocol as HEK293T. HskMC cells were cultured in Skeletal Muscle Cell Growth Medium, and transfected by using the same protocol as HEK293T as well.

### Small angle X-ray scattering (SAXS)

Synchrotron small-angle X-ray scattering (SAXS) measurements were conducted at the BL19U2 beamline of the Shanghai Synchrotron Radiation Facility (SSRF), utilizing an X-ray wavelength of 0.103 nm. The experimental setup included a Pilatus 2 M detector (172 μm × 172 μm) with a resolution of 1043 × 981 pixels. The sample-to-detector distance was maintained at 5.68 meters, with the scattering vector *q* spanning from 0.003 Å^-1^ to 0.208 Å^-1^. Data acquisition involved capturing twenty consecutive two-dimensional (2D) images, each with a 0.5-second exposure time. These 2D images were subsequently integrated into one-dimensional (1D) intensity profiles using BioXTAS RAW 2.3.0 software.^32^ The final intensity curves were obtained by averaging the data from all 20 frames and subtracting the corresponding solvent background. The SAXS characterization was performed at a fixed concentration of lipid of ∼1.3 mg/mL.

### Data analysis of SAXS

The scattering curves of a particle *I*(*q*) is a Fourier transform of its pair distance distribution function, *P*(*r*), being related to each other by the equation:^33^

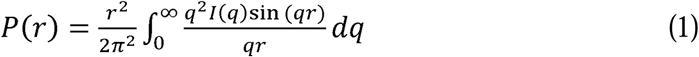

where *r* is the pair distance in real space, and *q* is the momentum transfer in reciprocal space. Furthermore, the radius of gyration (R_g_) can be calculated from the *P*(*r*) function using the equation:^33^

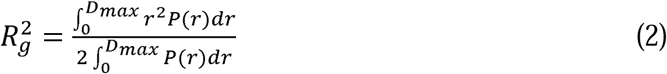

where *D_max_*is the maximum dimension of the particle.

SAXS data for mRNA-free LNPs was fitted using a broad peak model in SasView (https://www.sasview.org/) with details on the fitting constraints given in the SI. The SAXS data for mRNA-LNPs was fitted using a combination of the broad peak model and one Gaussian peak. The peak position from the broad peak model were fixed, with other parameters allowed to vary, while all of the Gaussian peak parameters were allowed to vary. For all samples, the data was fitted over the *q* range 0.02 – 0.2 Å^-1^. The areas of the broad peak and the Gaussian peak correspond to the contributions from disordered and ordered structures in the mRNA-LNP samples, respectively.

### Principal Component Analysis (PCA)

PCA was carried out to reduce dimensionality and visualize the structure of the dataset. All quantitative variables were first standardized to zero mean and unit variance. The PCA was performed with the PCA function from the FactoMineR package (2.12)^34^ in R (v 4.3.3). The resulting principal components were evaluated based on their eigenvalues and cumulative explained variance to determine the optimal number of components for downstream interpretation. Visualization of sample scores and variable loadings was generated using ggplot2.

### Nuclear Magnetic Resonance (NMR)

The ^31^P NMR spectra for concentrated mRNA, fresh mRNA-LNP, and lyophilized mRNA-LNP samples were acquired using a Bruker AVANCE NEO 700 MHz NMR spectrometer. This system was equipped with a 5 mm helium-cooled broadband cryo-probe and a wide-bore superconducting magnet providing a magnetic field strength of 16.4 T (700 MHz), located at Bruker BioSpin (Germany).

### Statistical analysis

To assess the significance of the observed differences among groups, Analysis of Variance (ANOVA) was conducted, followed by a Dunn-Sidak post-hoc test for multiple comparisons. A significance level of α = 0.05 was applied, and all pairwise comparisons were adjusted using the Dunn-Sidak correction within the statistical toolbox of Origin software.

## ASSOCIATED CONTENT

### Supporting Information

Cryo-EM images and SAXS curve of mRNA-LNPs predominantly exhibiting spherical morphology, stability tests for frozen and lyophilized mRNA-LNPs from 0 to 24 weeks, luminescence of mRNA-LNPs in HskMC, HepG2, HeLa, and HEK293T cells, all peaks present in the SAXS data for various mRNA-LNPs, detailed sample information for mRNA-LNPs, fitting method for SAXS data of mRNA-free LNPs and mRNA-LNPs, fitting parameters for the SAXS data of mRNA-free LNPs with the broad peak model, and fitting parameters of broad peak and Gaussian peak employed to fit the SAXS data for mRNA-LNPs.

## AUTHOR INFORMATION

### Author Contributions

Z.L. and L.H. designed and supervised the project. X.C., J.F., X.G., M.L., and F.J. prepared the samples and performed the measurements. X.C. and Z.L. analyzed the experimental data. X.C., J.F., X.G., Z.L., and L.H. wrote the manuscript. All authors have given approval to the final version of the manuscript.

## ACKNOWLEDGMENT

This study was supported by the Student Innovation Center at Shanghai Jiao Tong University. We thank the staff members of BL19U2 beamline (https://cstr.cn/31129.02.NFPS.BL19U2) at the National Facility for Protein Science in Shanghai (https://cstr.cn/31129.02.NFPS), for providing technical support and assistance in data collection and analysis.

## REFERENCES

1. Polack, F. P.; Thomas, S. J.; Kitchin, N.; Absalon, J.; Gurtman, A.; Lockhart, S.; Perez, J. L.; Pérez Marc, G.; Moreira, E. D.; Zerbini, C.; Bailey, R.; Swanson, K. A.; Roychoudhury, S.; Koury, K.; Li, P.; Kalina, W. V.; Cooper, D.; Frenck, R. W.; Hammitt, L. L.; Türeci, Ö.; Nell, H.; Schaefer, A.; Ünal, S.; Tresnan, D. B.; Mather, S.; Dormitzer, P. R.; Şahin, U.; Jansen, K. U.; Gruber, W. C., Safety and Efficacy of the BNT162b2 mRNA Covid-19 Vaccine. N. Engl. J. Med. 2020, 383 (27), 2603–2615.

2. Baden, L. R.; El Sahly, H. M.; Essink, B.; Kotloff, K.; Frey, S.; Novak, R.; Diemert, D.; Spector, S. A.; Rouphael, N.; Creech, C. B.; McGettigan, J.; Khetan, S.; Segall, N.; Solis, J.; Brosz, A.; Fierro, C.; Schwartz, H.; Neuzil, K.; Corey, L.; Gilbert, P.; Janes, H.; Follmann, D.; Marovich, M.; Mascola, J.; Polakowski, L.; Ledgerwood, J.; Graham, B. S.; Bennett, H.; Pajon, R.; Knightly, C.; Leav, B.; Deng, W.; Zhou, H.; Han, S.; Ivarsson, M.; Miller, J.; Zaks, T., Efficacy and Safety of the mRNA-1273 SARS-CoV-2 Vaccine. N. Engl. J. Med. 2020, 384 (5), 403–416.

3. Hou, X.; Zaks, T.; Langer, R.; Dong, Y., Lipid nanoparticles for mRNA delivery. Nat. Rev. Mater. 2021, 6 (12), 1078–1094.

4. Tenchov, R.; Bird, R.; Curtze, A. E.; Zhou, Q., Lipid Nanoparticles─From Liposomes to mRNA Vaccine Delivery, a Landscape of Research Diversity and Advancement. ACS Nano 2021, 15 (11), 16982–17015.

5. Barbier, A. J.; Jiang, A. Y.; Zhang, P.; Wooster, R.; Anderson, D. G., The clinical progress of mRNA vaccines and immunotherapies. Nat. Biotechnol. 2022, 40 (6), 840–854.

6. Weng, Y.; Li, C.; Yang, T.; Hu, B.; Zhang, M.; Guo, S.; Xiao, H.; Liang, X.-J.; Huang, Y., The challenge and prospect of mRNA therapeutics landscape. Biotechnol. Adv. 2020, 40, 107534.

7. Wilson, E.; Goswami, J.; Baqui Abdullah, H.; Doreski Pablo, A.; Perez-Marc, G.; Zaman, K.; Monroy, J.; Duncan Christopher, J. A.; Ujiie, M.; Rämet, M.; Pérez-Breva, L.; Falsey Ann, R.; Walsh Edward, E.; Dhar, R.; Wilson, L.; Du, J.; Ghaswalla, P.; Kapoor, A.; Lan, L.; Mehta, S.; Mithani, R.; Panozzo Catherine, A.; Simorellis Alana, K.; Kuter Barbara, J.; Schödel, F.; Huang, W.; Reuter, C.; Slobod, K.; Stoszek Sonia, K.; Shaw Christine, A.; Miller Jacqueline, M.; Das, R.; Chen Grace, L., Efficacy and Safety of an mRNA-Based RSV PreF Vaccine in Older Adults. N. Engl. J. Med. 2023, 389 (24), 2233–2244.

8. Cheng, M. H. Y.; Leung, J.; Zhang, Y.; Strong, C.; Basha, G.; Momeni, A.; Chen, Y.; Jan, E.; Abdolahzadeh, A.; Wang, X.; Kulkarni, J. A.; Witzigmann, D.; Cullis, P. R., Induction of Bleb Structures in Lipid Nanoparticle Formulations of mRNA Leads to Improved Transfection Potency. Adv. Mater. 2023, 35 (31), 2303370.

9. Patel, S.; Ashwanikumar, N.; Robinson, E.; Xia, Y.; Mihai, C.; Griffith, J. P.; Hou, S.; Esposito, A. A.; Ketova, T.; Welsher, K.; Joyal, J. L.; Almarsson, Ö.; Sahay, G., Naturally-occurring cholesterol analogues in lipid nanoparticles induce polymorphic shape and enhance intracellular delivery of mRNA. Nat. Commun. 2020, 11 (1), 983.

10. Yanez Arteta, M.; Kjellman, T.; Bartesaghi, S.; Wallin, S.; Wu, X.; Kvist, A. J.; Dabkowska, A.; Székely, N.; Radulescu, A.; Bergenholtz, J.; Lindfors, L., Successful reprogramming of cellular protein production through mRNA delivered by functionalized lipid nanoparticles. Proc. Natl. Acad. Sci. U. S. A. 2018, 115 (15), E3351–E3360.

11. Cui, L.; Hunter, M. R.; Sonzini, S.; Pereira, S.; Romanelli, S. M.; Liu, K.; Li, W.; Liang, L.; Yang, B.; Mahmoudi, N.; Desai, A. S., Mechanistic Studies of an Automated Lipid Nanoparticle Reveal Critical Pharmaceutical Properties Associated with Enhanced mRNA Functional Delivery In Vitro and In Vivo. Small 2022, 18 (9), 2105832.

12. Yu, H.; Iscaro, J.; Dyett, B.; Zhang, Y.; Seibt, S.; Martinez, N.; White, J.; Drummond, C. J.; Bozinovski, S.; Zhai, J., Inverse Cubic and Hexagonal Mesophase Evolution within Ionizable Lipid Nanoparticles Correlates with mRNA Transfection in Macrophages. J. Am. Chem. Soc. 2023, 145 (45), 24765–24774.

13. Yu, H.; Angelova, A.; Angelov, B.; Dyett, B.; Matthews, L.; Zhang, Y.; El Mohamad, M.; Cai, X.; Valimehr, S.; Drummond, C. J.; Zhai, J., Real-Time pH-Dependent Self-Assembly of Ionisable Lipids from COVID-19 Vaccines and In Situ Nucleic Acid Complexation. Angew. Chem. Int. Ed. 2023, 62 (35), e202304977.

14. Zheng, L.; Bandara, S. R.; Tan, Z.; Leal, C., Lipid nanoparticle topology regulates endosomal escape and delivery of RNA to the cytoplasm. Proc. Natl. Acad. Sci. U. S. A. 2023, 120 (27), e2301067120.

15. Philipp, J.; Dabkowska, A.; Reiser, A.; Frank, K.; Krzysztoń, R.; Brummer, C.; Nickel, B.; Blanchet, C. E.; Sudarsan, A.; Ibrahim, M.; Johansson, S.; Skantze, P.; Skantze, U.; Östman, S.; Johansson, M.; Henderson, N.; Elvevold, K.; Smedsrød, B.; Schwierz, N.; Lindfors, L.; Rädler, J. O., pH-dependent structural transitions in cationic ionizable lipid mesophases are critical for lipid nanoparticle function. Proc. Natl. Acad. Sci. U. S. A. 2023, 120 (50), e2310491120.

16. Sebastiani, F.; Yanez Arteta, M.; Lerche, M.; Porcar, L.; Lang, C.; Bragg, R. A.; Elmore, C. S.; Krishnamurthy, V. R.; Russell, R. A.; Darwish, T.; Pichler, H.; Waldie, S.; Moulin, M.; Haertlein, M.; Forsyth, V. T.; Lindfors, L.; Cárdenas, M., Apolipoprotein E Binding Drives Structural and Compositional Rearrangement of mRNA-Containing Lipid Nanoparticles. ACS Nano 2021, 15 (4), 6709–6722.

17. Brader, M. L.; Williams, S. J.; Banks, J. M.; Hui, W. H.; Zhou, Z. H.; Jin, L., Encapsulation state of messenger RNA inside lipid nanoparticles. Biophys. J. 2021, 120 (14), 2766–2770.

18. Gilbert, J.; Sebastiani, F.; Arteta, M. Y.; Terry, A.; Fornell, A.; Russell, R.; Mahmoudi, N.; Nylander, T., Evolution of the structure of lipid nanoparticles for nucleic acid delivery: From in situ studies of formulation to colloidal stability. J. Colloid Interface Sci. 2024, 660, 66–76.

19. Thelen, J. L.; Leite, W.; Urban, V. S.; O’Neill, H. M.; Grishaev, A. V.; Curtis, J. E.; Krueger, S.; Castellanos, M. M., Morphological Characterization of Self-Amplifying mRNA Lipid Nanoparticles. ACS Nano 2024, 18 (2), 1464–1476.

20. Li, Y.; Ma, C.; Han, Z.; Weng, W.; Yang, S.; He, Z.; Li, Z.; Su, X.; Zuo, T.; Cheng, H., Morphology evolution of lipid nanoparticle discovered by small angle neutron scattering. Giant 2024, 20, 100329.

21. Zhao, P.; Hou, X.; Yan, J.; Du, S.; Xue, Y.; Li, W.; Xiang, G.; Dong, Y., Long-term storage of lipid-like nanoparticles for mRNA delivery. Bioact. Mater. 2020, 5 (2), 358–363.

22. Fan, Y.; Rigas, D.; Kim, L. J.; Chang, F.-P.; Zang, N.; McKee, K.; Kemball, C. C.; Yu, Z.; Winkler, P.; Su, W.-C.; Jessen, P.; Hura, G. L.; Chen, T.; Koenig, S. G.; Nagapudi, K.; Leung, D.; Yen, C.-W., Physicochemical and structural insights into lyophilized mRNA-LNP from lyoprotectant and buffer screenings. J. Control. Release 2024, 373, 727–737.

23. Lamoot, A.; Lammens, J.; De Lombaerde, E.; Zhong, Z.; Gontsarik, M.; Chen, Y.; De Beer, T. R. M.; De Geest, B. G., Successful batch and continuous lyophilization of mRNA LNP formulations depend on cryoprotectants and ionizable lipids. Biomater. Sci. 2023, 11 (12), 4327–4334.

24. Li, M.; Jia, L.; Xie, Y.; Ma, W.; Yan, Z.; Liu, F.; Deng, J.; Zhu, A.; Siwei, X.; Su, W.; Liu, X.; Li, S.; Wang, H.; Yu, P.; Zhu, T., Lyophilization process optimization and molecular dynamics simulation of mRNA-LNPs for SARS-CoV-2 vaccine. npj Vaccines 2023, 8 (1), 153.

25. Kafetzis, K. N.; Papalamprou, N.; McNulty, E.; Thong, K. X.; Sato, Y.; Mironov, A.; Purohit, A.; Welsby, P. J.; Harashima, H.; Yu-Wai-Man, C.; Tagalakis, A. D., The Effect of Cryoprotectants and Storage Conditions on the Transfection Efficiency, Stability, and Safety of Lipid-Based Nanoparticles for mRNA and DNA Delivery. Adv. Healthc. Mater. 2023, 12 (18), 2203022.

26. Nogueira, S. S.; Schlegel, A.; Maxeiner, K.; Weber, B.; Barz, M.; Schroer, M. A.; Blanchet, C. E.; Svergun, D. I.; Ramishetti, S.; Peer, D.; Langguth, P.; Sahin, U.; Haas, H., Polysarcosine-Functionalized Lipid Nanoparticles for Therapeutic mRNA Delivery. ACS Appl. Nano Mater. 2020, 3 (11), 10634–10645.

27. Hassett, K. J.; Higgins, J.; Woods, A.; Levy, B.; Xia, Y.; Hsiao, C. J.; Acosta, E.; Almarsson, Ö.; Moore, M. J.; Brito, L. A., Impact of lipid nanoparticle size on mRNA vaccine immunogenicity. J. Control. Release 2021, 335, 237–246.

28. Leung, A. K. K.; Hafez, I. M.; Baoukina, S.; Belliveau, N. M.; Zhigaltsev, I. V.; Afshinmanesh, E.; Tieleman, D. P.; Hansen, C. L.; Hope, M. J.; Cullis, P. R., Lipid Nanoparticles Containing siRNA Synthesized by Microfluidic Mixing Exhibit an Electron-Dense Nanostructured Core. J. Phys. Chem. C 2012, 116 (34), 18440–18450.

29. DiVerdi, J. A.; Opella, S. J.; Ma, R. I.; Kallenbach, N. R.; Seeman, N. C., 31P NMR of DNA in eukaryotic chromosomal complexes. Biochem. Biophys. Res. Commun. 1981, 102 (3), 885–890.

30. Hammel, M.; Fan, Y.; Sarode, A.; Byrnes, A. E.; Zang, N.; Kou, P.; Nagapudi, K.; Leung, D.; Hoogenraad, C. C.; Chen, T.; Yen, C.-W.; Hura, G. L., Correlating the Structure and Gene Silencing Activity of Oligonucleotide-Loaded Lipid Nanoparticles Using Small-Angle X-ray Scattering. ACS Nano 2023, 17 (12), 11454–11465.

31. Padilla, M. S.; Shepherd, S. J.; Hanna, A. R.; Kurnik, M.; Zhang, X.; Chen, M.; Byrnes, J.; Joseph, R. A.; Yamagata, H. M.; Ricciardi, A. S.; Mrksich, K.; Issadore, D.; Gupta, K.; Mitchell, M. J., Elucidating lipid nanoparticle properties and structure through biophysical analyses. Nat. Biotechnol. 2025.

32. Hopkins, J., BioXTAS RAW 2: new developments for a free open-source program for small-angle scattering data reduction and analysis. J. Appl. Crystallogr. 2024, 57 (1), 194–208.

33. Dmitri, I. S.; Michel, H. J. K., Small-angle scattering studies of biological macromolecules in solution. Rep. Prog. Phys. 2003, 66 (10), 1735.

34. Lê, S.; Josse, J.; Husson, F., FactoMineR: An R Package for Multivariate Analysis. J. Stat. Softw. 2008, 25 (1), 1–18.

