## supporting information for "Revealing a Correlation between Structure and *in vitro* Activity of mRNA Lipid Nanoparticles"

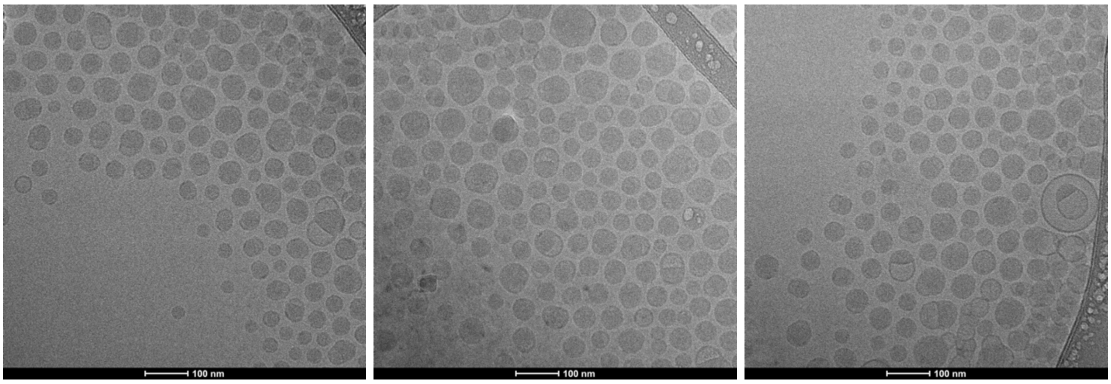


**Figure S1.** Cryo-EM image of mRNA-LNPs predominantly exhibiting spherical structures. The scale bar is 100 nm for each panel.

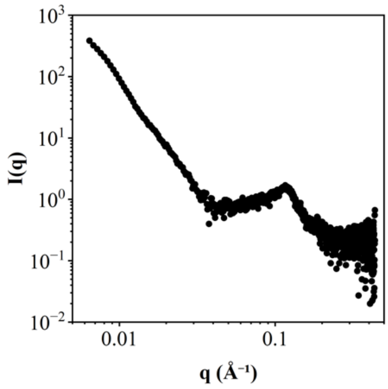


**Figure S2.** Small angle X-ray scattering curve of spherical mRNA-LNPs.

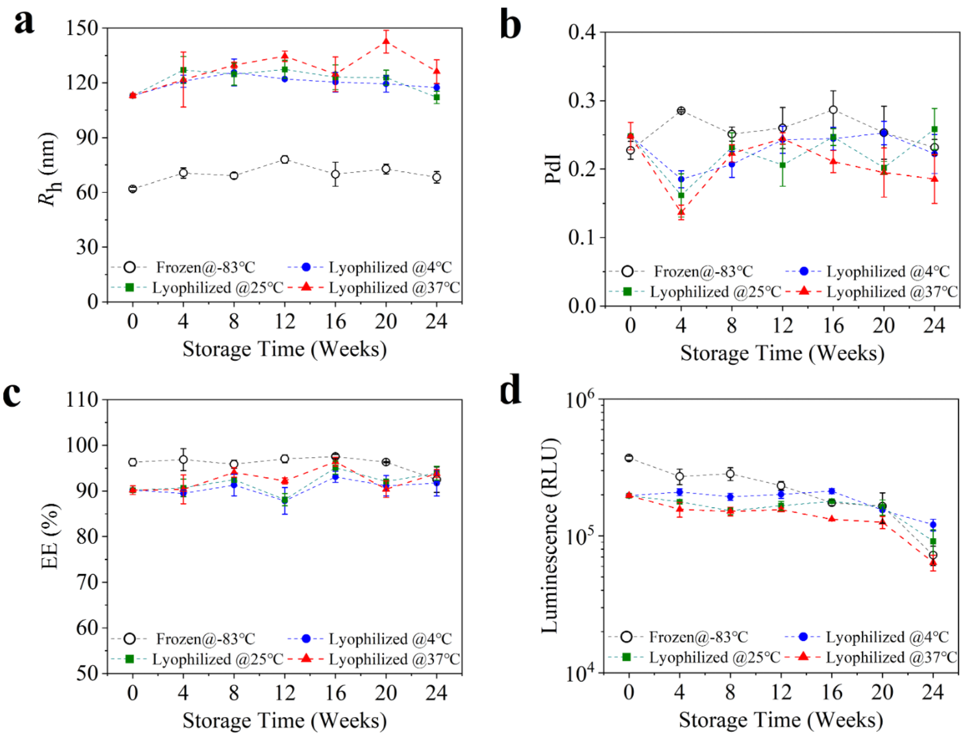


**Figure S3.** Stability tests for frozen mRNA-LNPs at -83 ℃ (black empty dots) and lyophilized mRNA-LNPs stored at 4 (blue solid dots), 25 (green solid squares), and 37 ℃ (red solid triangles) from 0 to 24 weeks. (a) Size, (b) Poly dispersive Index (PdI), (c) Encapsulation Efficiency (EE), and (d) Luminescence variations from 0 to 24 weeks.

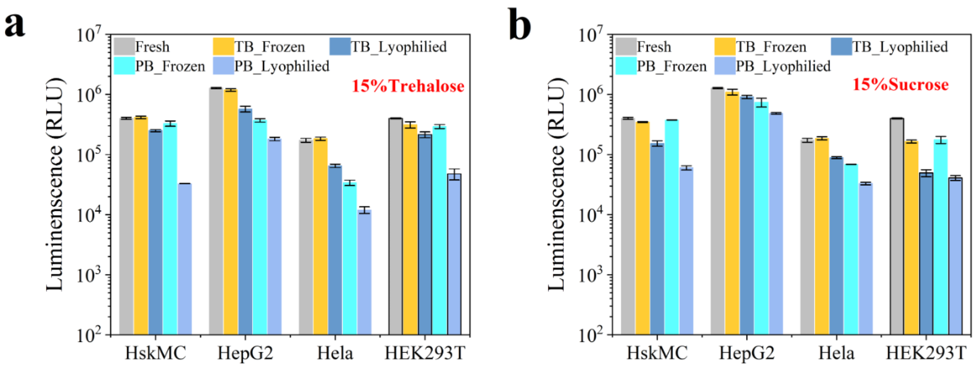


**Figure S4.** Luminescence of mRNA-LNPs in HskMC, HepG2, HeLa, and HEK293T cells. (a) Luminescence of fresh mRNA-LNPs (gray), and mRNA-LNPs frozen with 15% (w/v) trehalose in 40 mM Tris buffer (TB, yellow) or phosphate buffer (PB, cyan), and lyophilized with 15% (w/v) trehalose in 40 mM Tris buffer (TB, dark blue) or phosphate buffer (PB, light blue). (b) Luminescence of fresh mRNA-LNPs (gray), and mRNA-LNPs frozen with 15% (w/v) sucrose in 40 mM Tris buffer (TB, yellow) or phosphate buffer (PB, cyan), and lyophilized with 15% (w/v) sucrose in 40 mM Tris buffer (TB, dark blue) or phosphate buffer (PB, light blue). For fresh samples in the four cells, the Lumi data are the same with those in Figure 1c.

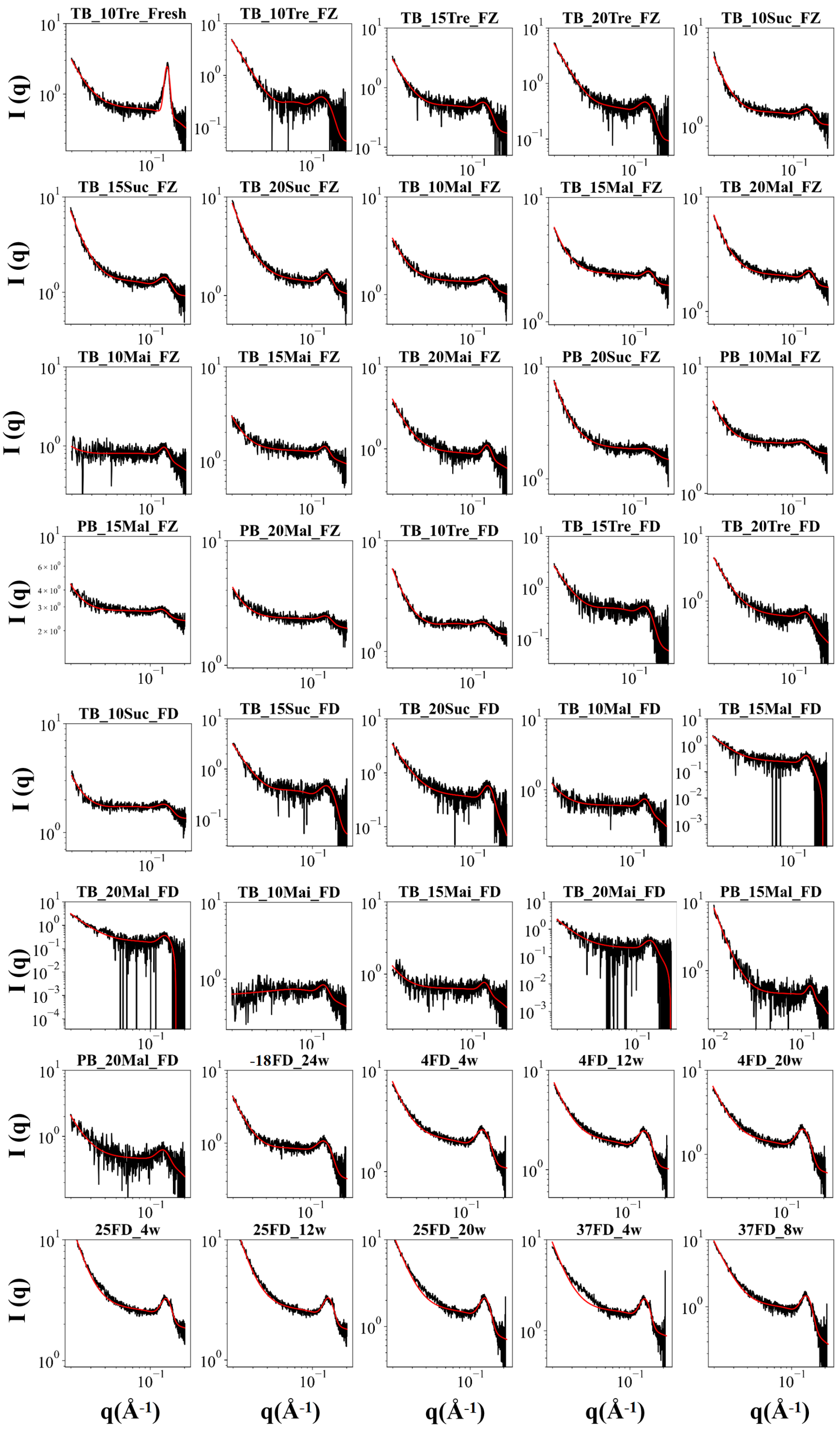


**Figure S5.** Representative peaks present in the SAXS data for various mRNA-LNPs were fitted with a broad peak model plus a Gaussian peak in a $q$ range from 0.02 Å$^{-1}$ to 0.2 Å$^{-1}$.

**Table S1.** Detailed sample information for mRNA-LNPs.

| Sample Name | Buffer | Lyoprotectant | Storage Temperature (℃) | Storage period (week) | Size (nm) | PdI | EE (%) |
|---|---|---|---|---|---|---|---|
| TB_10Tre_Fresh | TB | 10% Trehalose | / | / | 71.54±2.10 | 0.200±0.008 | 91.92±1.00 |
| TB_10Tre_FZ | TB | 10% Trehalose | -83 | 24 | 77.10±2.43 | 0.211±0.018 | 88.56±2.22 |
| TB_15Tre_FZ | TB | 15% Trehalose | -83 | 24 | 67.31±1.60 | 0.245±0.004 | 88.27±5.19 |
| TB_20Tre_FZ | TB | 20% Trehalose | -83 | 24 | 67.08±1.20 | 0.245±0.014 | 90.84±1.41 |
| TB_10Suc_FZ | TB | 10% Sucrose | -83 | 24 | 63.25±0.93 | 0.300±0.013 | 91.38±0.06 |
| TB_15Suc_FZ | TB | 15% Sucrose | -83 | 24 | 61.18±2.25 | 0.261±0.011 | 81.62±4.28 |
| TB_20Suc_FZ | TB | 20% Sucrose | -83 | 24 | 63.51±0.63 | 0.231±0.014 | 86.14±1.50 |
| TB_10Mal_FZ | TB | 10% Maltose | -83 | 24 | 87.67±1.00 | 0.215±0.029 | 91.57±2.46 |
| TB_15Mal_FZ | TB | 15% Maltose | -83 | 24 | 85.98±5.28 | 0.248±0.055 | 66.04±5.20 |
| TB_20Mal_FZ | TB | 20% Maltose | -83 | 24 | 86.09±2.02 | 0.286±0.012 | 93.36±1.35 |
| TB_10Mai_FZ | TB | 10% Maltitol | -83 | 24 | 101.59±1.49 | 0.200±0.025 | 94.51±0.75 |
| TB_15Mai_FZ | TB | 15% Maltitol | -83 | 24 | 109.38±3.28 | 0.255±0.010 | 86.35±9.22 |
| TB_20Mai_FZ | TB | 20% Maltitol | -83 | 24 | 109.16±3.32 | 0.215±0.032 | 89.54±0.16 |
| PB_10Tre_FZ | PB | 10% Trehalose | -83 | 24 | 123.06±1.78 | 0.224±0.035 | 86.35±2.92 |
| PB_15Tre_FZ | PB | 15% Trehalose | -83 | 24 | 116.67±3.37 | 0.213±0.019 | 87.72±6.68 |
| PB_20Tre_FZ | PB | 20% Trehalose | -83 | 24 | 107.48±0.42 | 0.220±0.026 | 88.24±4.36 |
| PB_10Suc_FZ | PB | 10% Sucrose | -83 | 24 | 98.33±0.82 | 0.167±0.042 | 80.72±1.15 |
| PB_15Suc_FZ | PB | 15% Sucrose | -83 | 24 | 93.35±2.02 | 0.164±0.031 | 93.97±0.69 |
| PB_20Suc_FZ | PB | 20% Sucrose | -83 | 24 | 87.62±0.65 | 0.188±0.018 | 89.57±0.06 |
| PB_10Mal_FZ | PB | 10% Maltose | -83 | 24 | 133.32±1.11 | 0.234±0.020 | 89.03±2.40 |
| PB_15Mal_FZ | PB | 15% Maltose | -83 | 24 | 126.15±3.08 | 0.256±0.021 | 93.64±4.59 |
| PB_20Mal_FZ | PB | 20% Maltose | -83 | 24 | 120.23±2.06 | 0.231±0.028 | 84.31±2.28 |
| PB_10Mai_FZ | PB | 10% Maltitol | -83 | 24 | 152.90±2.72 | 0.209±0.022 | 86.23±1.02 |
| PB_15Mai_FZ | PB | 15% Maltitol | -83 | 24 | 161.60±1.97 | 0.244±0.011 | 88.73±2.03 |
| PB_20Mai_FZ | PB | 20% Maltitol | -83 | 24 | 186.80±9.60 | 0.260±0.023 | 67.22±4.37 |
| TB_10Tre_FD | TB | 10% Trehalose | / | / | 105.99±2.65 | 0.197±0.054 | 91.42±1.00 |
| TB_15Tre_FD | TB | 15% Trehalose | / | / | 104.97±2.10 | 0.204±0.036 | 91.38±1.12 |
| TB_20Tre_FD | TB | 20% Trehalose | / | / | 297.42±26.66 | 0.232±0.053 | 90.20±0.38 |
| TB_10Suc_FD | TB | 10% Sucrose | / | / | 95.12±5.90 | 0.186±0.024 | 90.09±3.13 |
| TB_15Suc_FD | TB | 15% Sucrose | / | / | 83.52±2.20 | 0.232±0.030 | 87.77±2.77 |
| TB_20Suc_FD | TB | 20% Sucrose | / | / | 91.15±2.94 | 0.240±0.018 | 89.68±1.40 |
| TB_10Mal_FD | TB | 10% Maltose | / | / | 151.34±15.32 | 0.239±0.044 | 85.88±6.44 |
| TB_15Mal_FD | TB | 15% Maltose | / | / | 215.00±27.84 | 0.212±0.034 | 87.33±1.08 |
| TB_20Mal_FD | TB | 20% Maltose | / | / | 154.33±21.43 | 0.288±0.021 | 87.48±3.91 |
| TB_10Mai_FD | TB | 10% Maltitol | / | / | 261.39±14.38 | 0.248±0.061 | 85.00±3.16 |
| TB_15Mai_FD | TB | 15% Maltitol | / | / | 211.97±19.43 | 0.272±0.036 | 84.72±3.29 |
| TB_20Mai_FD | TB | 20% Maltitol | / | / | 254.26±19.48 | 0.239±0.041 | 86.44±1.56 |
| PB_10Tre_FD | PB | 10% Trehalose | / | / | 291.19±17.38 | 0.317±0.023 | 62.39±5.74 |
| PB_15Tre_FD | PB | 15% Trehalose | / | / | 292.10±30.25 | 0.269±0.042 | 74.53±4.66 |
| PB_20Tre_FD | PB | 20% Trehalose | / | / | 497.20±41.01 | 0.316±0.035 | 84.16±8.16 |
| PB_10Suc_FD | PB | 10% Sucrose | / | / | 172.11±5.22 | 0.265±0.055 | 49.05±5.81 |
| PB_15Suc_FD | PB | 15% Sucrose | / | / | 165.93±4.63 | 0.241±0.008 | 69.19±6.21 |
| PB_20Suc_FD | PB | 20% Sucrose | / | / | 185.42±6.67 | 0.239±0.021 | 82.76±2.27 |
| PB_10Mal_FD | PB | 10% Maltose | / | / | 431.69±22.83 | 0.321±0.040 | 81.84±2.36 |

| | | | | | | | |
|---|---|---|---|---|---|---|---|
| PB_15Mal_FD | PB | 15% Maltose | / | / | 361.41±59.85 | 0.270±0.024 | 89.48±3.45 |
| PB_20Mal_FD | PB | 20% Maltose | / | / | 236.87±9.07 | 0.281±0.019 | 91.62±3.19 |
| PB_10Mai_FD | PB | 10% Maltitol | / | / | 271.61±21.22 | 0.266±0.005 | 51.83±8.23 |
| PB_15Mai_FD | PB | 15% Maltitol | / | / | 287.22±13.98 | 0.268±0.035 | 44.61±3.55 |
| PB_20Mai_FD | PB | 20% Maltitol | / | / | 391.08±85.39 | 0.274±0.017 | 43.30±5.33 |
| -18FD_24w | TB | 15% Trehalose | -18 | 24 | 68.25±3.11 | 0.232±0.012 | 92.48±2.79 |
| 4FD_4w | TB | 15% Trehalose | 4 | 4 | 120.91±3.32 | 0.185±0.012 | 94.89±1.19 |
| 4FD_12w | TB | 15% Trehalose | 4 | 12 | 122.07±0.27 | 0.243±0.020 | 92.55±0.09 |
| 4FD_20w | TB | 15% Trehalose | 4 | 20 | 119.43±4.48 | 0.253±0.017 | 93.44±1.78 |
| 25FD_4w | TB | 15% Trehalose | 25 | 4 | 127.11±7.40 | 0.162±0.032 | 92.85±2.45 |
| 25FD_12w | TB | 15% Trehalose | 25 | 12 | 127.37±4.99 | 0.206±0.031 | 91.25±3.25 |
| 25FD_20w | TB | 15% Trehalose | 25 | 20 | 122.86±4.08 | 0.202±0.009 | 93.18±2.52 |
| 37FD_4w | TB | 15% Trehalose | 37 | 4 | 121.81±5.06 | 0.137±0.011 | 94.33±1.48 |
| 37FD_8w | TB | 15% Trehalose | 37 | 8 | 129.73±1.58 | 0.223±0.013 | 91.57±0.43 |

TB and PB correspond to Tris and phosphate buffer, respectively.

**Data fitting for SAXS measurements of drug-free LNPs and mRNA-LNPs.**

The SAXS data for drug-free LNPs were fitted using the broad peak model implemented in SASView (https://www.sasview.org/), with all fitting parameters permitted to vary. The resulting parameters are detailed in Table S2. For the mRNA-LNPs, the SAXS data were fitted using a combination of the broad peak model and a Gaussian peak. The broad peak position and Porod exponent were fixed based on the drug-free LNP fitting results, while the other parameters were variable, along with all parameters of the Gaussian peak. Figure S6 illustrates the fitting curves for representative mRNA-LNP samples, with the corresponding fitting parameters provided in Table S3.

**Table S2.** Fitting parameters for the SAXS data of drug-free LNPs with the broad peak model.

| Parameters | Value |
|---|---|
| Porod scale | $1.88 \times 10^{-5}$ |
| Porod exponent | 3.696 |
| Lorentz scale | 2.647 |
| Lorentz length | 13.211 |
| Peak position | 0.06964 |

**Table S3.** Parameters of broad peak and Gaussian peak employed to fit the SAXS data for mRNA-LNPs.

| Sample name | Board peak | | Gaussian Peak | | |
|---|---|---|---|---|---|
| | Porod scale | Lorentz scale | Scale | Peak center | σ |
| TB_10Tre_Fresh | 1.312E-06 | 0.3016 | 1.9436 | 0.1382 | 6.592E-03 |
| TB_10Tre_FZ | 2.480E-06 | 0.1886 | 0.3078 | 0.1234 | 2.272E-02 |
| TB_15Tre_FZ | 1.399E-06 | 0.3451 | 0.2990 | 0.1276 | 1.782E-02 |
| TB_20Tre_FZ | 2.600E-06 | 0.2728 | 0.3156 | 0.1276 | 1.951E-02 |
| TB_10Suc_FZ | 2.044E-06 | 0.3249 | 0.3972 | 0.1314 | 1.855E-02 |
| TB_15Suc_FZ | 3.163E-06 | 0.3641 | 0.4383 | 0.1339 | 1.801E-02 |
| TB_20Suc_FZ | 3.818E-06 | 0.3826 | 0.4419 | 0.1356 | 1.475E-02 |
| TB_10Mal_FZ | 1.308E-06 | 0.3668 | 0.2508 | 0.1342 | 1.590E-02 |
| TB_15Mal_FZ | 1.845E-06 | 0.4958 | 0.4166 | 0.1361 | 1.397E-02 |
| TB_20Mal_FZ | 2.638E-06 | 0.5347 | 0.4536 | 0.1374 | 1.356E-02 |
| TB_10Mai_FZ | 1.255E-07 | 0.5000 | 0.2514 | 0.1307 | 1.107E-02 |
| TB_15Mai_FZ | 9.352E-07 | 0.5000 | 0.2699 | 0.1319 | 1.133E-02 |
| TB_20Mai_FZ | 1.680E-06 | 0.5000 | 0.3286 | 0.1359 | 1.071E-02 |
| PB_20Suc_FZ | 2.969E-06 | 0.5000 | 0.1509 | 0.1355 | 6.592E-03 |
| PB_10Mal_FZ | 1.584E-06 | 0.5436 | 0.2457 | 0.1239 | 1.827E-02 |
| PB_15Mal_FZ | 9.347E-07 | 0.4828 | 0.2727 | 0.1280 | 1.619E-02 |
| PB_20Mal_FZ | 1.026E-06 | 0.4463 | 0.2815 | 0.1323 | 1.508E-02 |
| TB_10Tre_FD | 1.762E-07 | 0.3561 | 0.9815 | 0.1357 | 1.499E-02 |
| TB_15Tre_FD | 9.288E-07 | 0.9413 | 1.6314 | 0.1376 | 5.004E-03 |
| TB_20Tre_FD | 2.181E-06 | 0.5000 | 0.2110 | 0.1295 | 6.541E-03 |
| TB_10Suc_FD | 9.606E-07 | 0.3816 | 0.3721 | 0.1320 | 1.686E-02 |
| TB_15Suc_FD | 1.564E-06 | 0.1766 | 0.3542 | 0.1344 | 1.531E-02 |
| TB_20Suc_FD | 1.586E-06 | 0.2646 | 0.3190 | 0.1387 | 2.006E-02 |
| TB_10Mal_FD | 3.490E-07 | 0.5952 | 0.2260 | 0.1307 | 2.006E-02 |
| TB_15Mal_FD | 1.014E-06 | 0.2318 | 0.2511 | 0.1330 | 1.961E-02 |
| TB_20Mal_FD | 1.555E-06 | 0.1900 | 0.2747 | 0.1359 | 1.311E-02 |
| TB_10Mai_FD | 3.311E-73 | 0.7392 | 0.2300 | 0.1287 | 1.227E-02 |
| TB_15Mai_FD | 3.771E-07 | 0.6249 | 0.2122 | 0.1289 | 1.127E-02 |
| TB_20Mai_FD | 1.090E-06 | 0.2036 | 0.2310 | 0.1313 | 1.397E-02 |
| PB_15Mal_FD | 3.171E-07 | 0.3605 | 0.2018 | 0.1268 | 1.149E-02 |
| PB_20Mal_FD | 8.572E-07 | 0.5043 | 0.2109 | 0.1300 | 1.514E-02 |
| -18FD_24w | 1.999E-06 | 0.5142 | 0.5964 | 0.1339 | 2.030E-02 |
| 4FD_4w | 8.137E-06 | 0.9352 | 1.2282 | 0.1378 | 1.764E-02 |
| 4FD_12w | 7.925E-06 | 0.8720 | 1.1743 | 0.1375 | 1.721E-02 |
| 4FD_20w | 7.113E-06 | 0.7868 | 1.1146 | 0.1371 | 1.708E-02 |
| 25FD_4w | 6.282E-06 | 0.8659 | 1.0947 | 0.1383 | 1.412E-02 |
| 25FD_12w | 7.203E-06 | 0.9030 | 1.1641 | 0.1386 | 1.387E-02 |
| 25FD_20w | 5.680E-06 | 0.8414 | 1.0941 | 0.1364 | 1.483E-02 |
| 37FD_4w | 4.235E-06 | 0.8201 | 1.0496 | 0.1368 | 1.429E-02 |
| 37FD_8w | 4.645E-06 | 0.7741 | 0.8774 | 0.1365 | 1.494E-02 |
